# ATG16L2 creates a VAIL-dedicated heterodimer with ATG16L1

**DOI:** 10.64898/2026.06.01.729256

**Authors:** Håvard Styrkestad Haukaas, Melanie Schwach, Laura Rodriguez de la Ballina, Namrita Kaur, Ann-Christin Borchers, Sven R Carlsson, Harald Stenmark, Alf Håkon Lystad

## Abstract

The ATG8 conjugation machinery supports both canonical autophagy, in which ATG8 proteins are lipidated on forming autophagosomes, and CASM, in which ATG8 proteins are conjugated to stressed single-membrane compartments. How cells allocate this shared machinery between these competing membrane programs remains unclear. Here, we identify ATG16L2 as a specialized regulator of V-ATPase-responsive ATG8 lipidation (atg8ylation). ATG16L2 does not form stable homodimers and is inactive in the absence of ATG16L1, but it becomes functional through heterodimerization with ATG16L1. The ATG16L1-ATG16L2 heterodimer constitutes a VAIL-competent E3-like complex that is excluded from WIPI2-positive autophagic membranes but readily recruited to stressed lysosomes. ATG16L2 also confers VAIL activity onto the otherwise VAIL-deficient ATG16L1α isoform. We propose that ATG16L1 homodimers and ATG16L1–ATG16L2 heterodimers represent alternative assemblies of the ATG8 conjugation machinery: ATG16L1α homodimers are dedicated to WIPI2-positive autophagic membranes, whereas ATG16L1–ATG16L2 heterodimers containing either ATG16L1 isoform form a VAIL-selective assembly.

## Introduction

Conjugation of mammalian homologues of yeast Atg8 to membrane lipids is a hallmark of both macroautophagy and CASM (conjugation of ATG8 to single membranes) (Hooper *et al*, 2022; Kabeya *et al*, 2004). These mammalian Atg8 homologues comprise the LC3 and GABARAP subfamilies, and we refer to their membrane conjugation collectively as ATG8 lipidation or atg8ylation. In macroautophagy, atg8ylation accompanies the *de novo* formation of double-membrane autophagosomes. In CASM, by contrast, LC3 and GABARAP proteins are conjugated to pre-existing single-membrane compartments, including lysosomes, endosomes and the Golgi, in response to membrane damage or other stress signals (Durgan & Florey, 2022).

Targeting of the ATG8 conjugation machinery to these distinct membranes is mediated by two E3-like complexes: ATG12-ATG5-ATG16L1 and ATG12-ATG5-TECPR1 (Boyle *et al*, 2023; Corkery *et al*, 2023; Fujita *et al*, 2008; Hanada *et al*, 2007; Kaur *et al*, 2023). During canonical autophagy, the ATG16L1 complex is recruited to PtdIns(3)P-positive phagophores through WIPI2 binding to ATG16L1 (Dooley *et al*, 2014; Gong *et al*, 2023). By contrast, CASM proceeds through two distinct branches: the sphingomyelin–TECPR1-induced LC3 lipidation pathway, STIL, which is triggered by membrane damage to endolysosomal compartments, and the V-ATPase–ATG16L1-induced LC3 lipidation pathway, VAIL, which is activated by disruption of the proton gradient across acidified membrane compartments (Figueras-Novoa *et al*, 2024).

Unlike TECPR1, which functions selectively in CASM, ATG16L1 operates in both canonical autophagy and CASM. This dual functionality is shaped by the two ATG16L1 splice variants, ATG16L1α and ATG16L1β, both of which support canonical autophagy but differ in VAIL competence. During CASM-inducing stress, ATG16L1β is efficiently recruited to stressed single-membrane compartments and supports robust VAIL, whereas ATG16L1α remains largely confined to WIPI2-positive autophagic structures (Lystad *et al*, 2019). This shows that a pool of ATG16L1α preserves autophagy during CASM-inducing stress by remaining in an autophagy-committed homodimeric state at WIPI2-positive membranes. How cells at the same time sustain a dedicated VAIL-active pool has remained unclear.

ATG16L2 is a close paralogue of ATG16L1 and shares its overall domain architecture: an N-terminal ATG5-interaction motif (AFIM), a central coiled-coil domain (CCD) that mediates dimerization, and a C-terminal seven-bladed WD40 β-propeller (**Fig. 1A**) (Ishibashi *et al*, 2011; Kim *et al*, 2015). Similar to ATG16L1, ATG16L2 also exists as multiple splice isoforms. In mice, two Atg16L2 splice variants have been described: a longer isoform containing all annotated exons and a shorter isoform lacking exon 8 (Ishibashi *et al*., 2011). As the full-length ATG16L2 isoform was reported to predominate in most mouse tissues, we focus on this isoform here.

**Figure 1.**
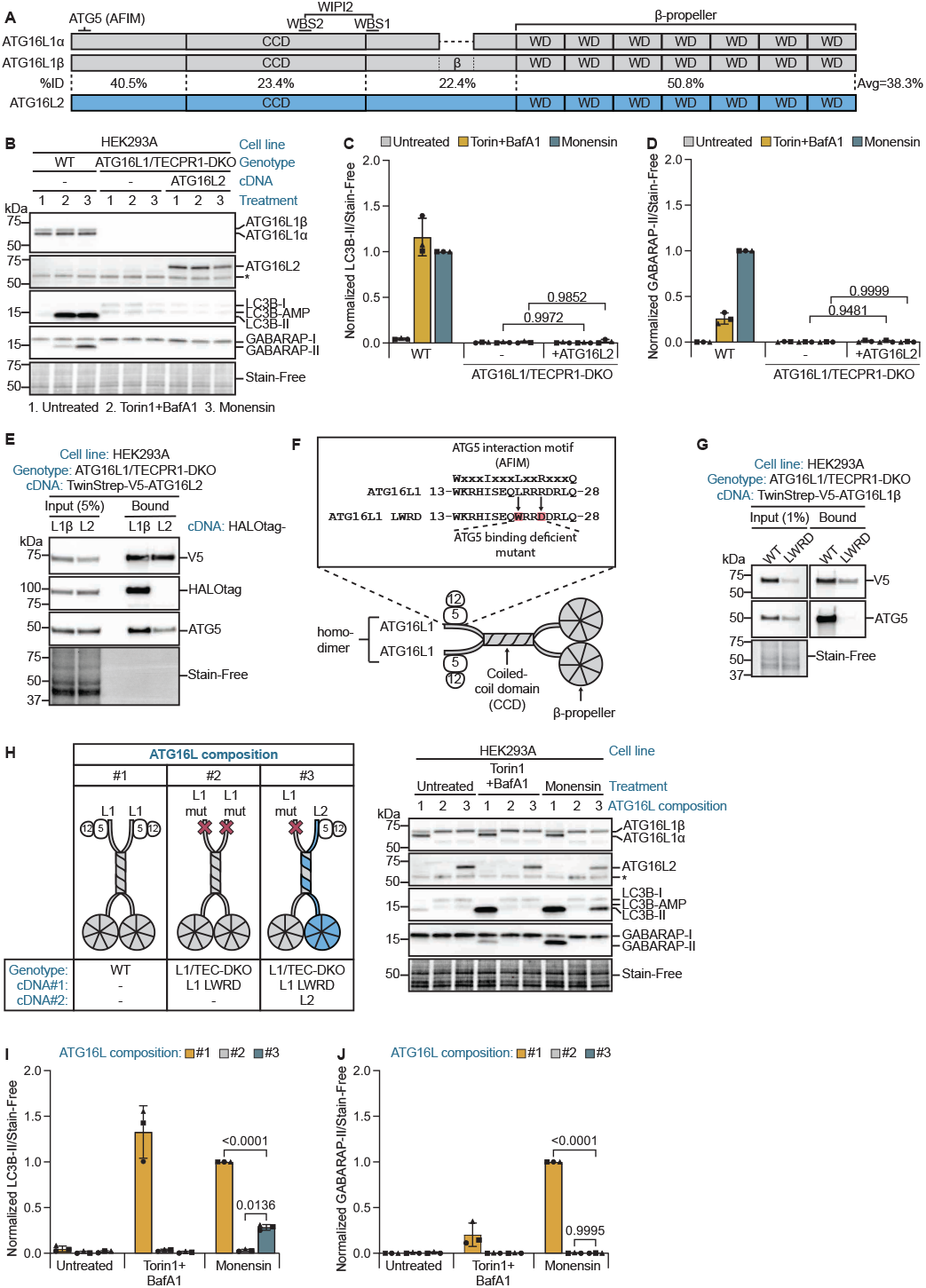
ATG16L2 is a specialized CASM regulator that requires heterodimerization with ATG16L1. **A**. Schematic overview of ATG16 family proteins. ATG16L1α, ATG16L1β and ATG16L2 share an N-terminal ATG5-interacting motif (AFIM), a central coiled-coil domain (CCD) and a C-terminal WD40 β-propeller. The β isoform-specific insertion in ATG16L1β and the WIPI2-interaction region present in ATG16L1 but absent from ATG16L2 are indicated. Percent amino-acid identity between ATG16L1β and ATG16L2 is shown for the N-terminal region, CCD, middle/disordered region and WD40 β-propeller, highlighting strong conservation of the N-terminal AFIM-containing region and WD40 β-propeller, but limited conservation within the CCD and intervening region. **B**. Immunoblot analysis of LC3B and GABARAP lipidation in wild-type (WT) and ATG16L1/TECPR1-DKO HEK293A cells with or without ATG16L2 reconstitution, left untreated (1) or treated with 50 nM Torin1 and 100 nM bafilomycin A1 (2) or 5 µM monensin (3) for 2 h. Stain-Free is a loading control. **C, D**. Quantification of LC3B-II (**C**) and GABARAP-II (**D**) from immunoblots in (**B**), normalized to Stain-Free. **E**. Strep-TactinXT pull-down of TwinStrep-V5-ATG16L2 from ATG16L1/TECPR1-DKO HEK293A cells co-expressing HaloTag-ATG16L1β or HaloTag-ATG16L2. Input and pull-down fractions were immunoblotted for the indicated proteins. **F**. Schematic of the ATG16L1 AFIM mutant (L21W/R24D; LWRD) used to disrupt ATG5 binding. **G**. Strep-TactinXT pull-down of TwinStrep-V5-ATG16L1β WT or LWRD mutant from ATG16L1/TECPR1-DKO HEK293A cells. Input and pull-down fractions were immunoblotted for the indicated proteins. **H**. Schematic of the ATG16L compositions analyzed and corresponding immunoblot of LC3B and GABARAP lipidation in HEK293A cells expressing the indicated ATG16L combinations, left untreated or treated with 50 nM Torin1 plus 100 nM bafilomycin A1 or 5 µM monensin for 2 h. ATG16L composition #1, WT cells expressing endogenous ATG16L1; ATG16L composition #2, ATG16L1/TECPR1-DKO cells expressing ATG16L1β L21W/R24D; ATG16L composition #3, ATG16L1/TECPR1-DKO cells co-expressing ATG16L1β L21W/R24D and ATG16L2. Stain-Free is a loading control. **I, J**. Quantification of LC3B-II (**I**) and GABARAP-II (**J**) from immunoblots in (**H**), normalized to Stain-Free. **Data information:** Quantifications are normalized as indicated in the graphs. Bars show mean ± SD; symbols denote independent experiments. Statistical significance was determined by two-way ANOVA followed by Tukey’s multiple-comparisons test (**C,D,I,J**). Adjusted p-values are indicated in the graphs.

Although ATG16L1β and ATG16L2 share the same overall architecture, they show limited sequence conservation, with 38.5% identity overall and only 23.4% identity within the CCD. This divergence is functionally relevant because WIPI2 binds ATG16L1 through two adjacent WIPI2-binding sites, WBS1 and WBS2. WBS1 lies adjacent to the CCD, whereas WBS2 is located within the CCD at the dimer interface. Both sites are present in ATG16L1α and ATG16L1β but not in ATG16L2 (Dooley *et al*., 2014; Gong *et al*., 2023). Consistent with the loss of these WIPI2-binding determinants, earlier work showed that ATG16L2 does not support canonical autophagy and identified the CCD as a key determinant of this functional difference (Ishibashi *et al*., 2011).

Here, we identify ATG16L2 as a functional counterpart to ATG16L1α in the balance between autophagy and CASM. Whereas ATG16L1α remains associated with WIPI2-positive autophagic membranes and thereby preserves an autophagy-committed ATG16L1 pool, ATG16L2 redirects either ATG16L1 isoform into a VAIL-responsive complex that is excluded from WIPI2-positive structures and recruited to stressed lysosomes. Although ATG16L2 is inactive on its own, heterodimerization with ATG16L1α or ATG16L1β unlocks its activity, expands the pool of VAIL-exclusive ATG16 complexes, and confers VAIL competence on the otherwise VAIL-deficient ATG16L1α isoform. These findings identify ATG16L1– ATG16L2 heterodimers as a dedicated V-ATPase-responsive branch of the ATG8 conjugation machinery.

## Results

### ATG16L2 is a specialized CASM regulator that requires heterodimerization with ATG16L1

The physiological role of ATG16L2 has remained unclear, perhaps in part because it is expressed at negligible levels in most commonly used cell lines. Consistent with Human Protein Atlas data (Jin *et al*, 2023), we were unable to detect ATG16L2 in HEK293A cells by MS and observed detectable ATG16L2 peptides in only one of five U2OS samples (**Figure EV1A**). This allowed us to use these cell lines as effectively zero-background systems for reconstitution of exogenous ATG16L2. To test whether ATG16L2 can support atg8ylation (ATG8 lipidation), we expressed ATG16L2 in ATG16L1/TECPR1 double-knockout (DKO) HEK293A cells, which are devoid of atg8ylation. Under both canonical autophagy-inducing conditions (Torin1) and CASM-inducing conditions (monensin), ATG16L2 failed to restore detectable LC3B or GABARAP lipidation (**Fig. 1B–D**).

Because ATG16 family proteins dimerize through their CCDs, we hypothesized that ATG16L2 might function as a latent binding partner that becomes active only upon heterodimerization with ATG16L1. To test its ability to form homo- and heterodimers, we expressed TwinStrep-V5-tagged ATG16L2 in ATG16L1/TECPR1-DKO HEK293A cells and complemented these cells with either HaloTag-ATG16L1β or HaloTag-ATG16L2 (**Fig. 1E**). TwinStrep-V5-ATG16L2 robustly co-precipitated HaloTag-ATG16L1β, whereas HaloTag-ATG16L2 was not recovered, indicating that ATG16L2 readily forms stable heterodimers with ATG16L1β but does not efficiently form stable homodimers. This preference for heterodimerization was not altered in response to monensin treatment (**Figure EV1C**). Notably,ATG16L2 alone still co-precipitated ATG5, indicating that it can engage ATG5 without detectable homodimer formation, although less efficiently than the ATG16L1-ATG16L2 heterodimer, consistent with the presence of only a single ATG5-binding site.

We next asked whether the ATG16L1β-ATG16L2 heterodimer is catalytically competent. Because both wild-type ATG16L1β and ATG16L2 contain an N-terminal ATG5-interaction motif, the native heterodimer is predicted to contain two potential ATG5-binding arms. To enforce dependence on the ATG16L2 arm, we generated an ATG16L1β mutant defective in ATG5 binding by introducing L21W and R24D substitutions into the AFIM (Kim *et al*., 2015). We then established a complementation system in ATG16L1/TECPR1-DKO cells co-expressing wild-type ATG16L2 together with AFIM-mutant ATG16L1β, thereby converting the heterodimer into a single-ATG5-binding-arm complex. In this system, the L21W/R24D substitutions abolished ATG5 binding and reduced ATG16L1β stability (**Fig. 1F,G,H; Figure EV1B**); however, mutant protein levels remained near endogenous levels of ATG16L1 (**Fig. 1H**).

Whereas the ATG16L1β mutant alone was inactive, co-expression with ATG16L2 restored LC3B lipidation selectively in response to monensin (**Fig. 1H–J**). This rescue was partial, relative to wild-type cells, consistent with ATG16L2 supplying the only functional ATG5-binding interface in this engineered heterodimer and therefore operating with reduced efficiency. We obtained the same result using an independent ATG16L1 mutant lacking the N-terminal 69 amino acids, which removes the entire ATG5-interaction region, further supporting that ATG16L2 can provide the functional ATG5-binding arm within the active heterodimer (**Figure EV1D,E**).

The recovery of activity from two individually inactive proteins identifies the ATG16L1β-ATG16L2 heterodimer as a functional E3-like complex. Conversely, the inability of ATG16L2 to support atg8ylation in ATG16L1-deficient cells is consistent with its apparent inability to form stable homodimers. Notably, heterodimer activity was not induced by Torin1-mediated autophagy, but neither was it restricted to monensin. Instead, it was triggered by a broader panel of CASM-inducing stimuli, including ionophore stress and lysosomal damage, closely mirroring the activation profile of the canonical VAIL pathway (**Figure EV1F–M**). Interestingly, in this single-ATG5-binding-arm configuration, the heterodimer supported LC3B but not GABARAP lipidation. This suggests either that efficient GABARAP conjugation requires both ATG5-binding interfaces within the dimer, or that ATG16L2 is intrinsically unable to support GABARAP lipidation; this is resolved below using chimeric constructs (Results section 3) and the native heterodimer (Result section 5).

### ATG16L2 is excluded from WIPI2 structures and recruited to compromised lysosomes

Before examining ATG16L2 localization by microscopy, we tested whether ATG16L1–ATG16L2-dependent CASM could be reproduced in U2OS cells using untagged proteins. As in HEK293A cells, expression of untagged ATG16L1β L21W/R24D alone did not restore LC3B lipidation in ATG16L1/TECPR1-DKO U2OS cells, whereas co-expression with untagged ATG16L2 rescued monensin-induced LC3B lipidation (**Figure EV2A,B**). Having confirmed ATG16L1β–ATG16L2-dependent CASM activity in this cell type, we next generated ATG16L1/TECPR1-DKO U2OS cells reconstituted with HaloTag-ATG16L1β and mNeonGreen-ATG16L2 to examine heterodimer localization. Treatment with Torin1 and bafilomycin A1 increased the number of ATG16L1β-positive WIPI2 structures, but these structures lacked ATG16L2 (**Fig. 2A,B**), indicating that ATG16L1β–ATG16L2 heterodimers are not recruited to sites of canonical autophagy. As expected, neither ATG16L1β nor ATG16L2 localized to LAMP1-positive lysosomes under Torin1 plus bafilomycin A1 treatment, confirming that lysosomal recruitment of these proteins is specific to CASM. In contrast, monensin, which induces both autophagy and CASM, recruited ATG16L1β to both WIPI2- and LAMP1-positive structures (**Fig. 2A–D**). Importantly, ATG16L2 remained absent from monensin-induced WIPI2 structures but was robustly recruited to LAMP1-positive compartments. Together, these data show that ATG16L2-containing heterodimers are directed to stressed lysosomes rather than WIPI2-positive autophagic membranes, and as such dedicate a pool of ATG16L1 containing complexes in the cell toward VAIL.

**Figure 2.**
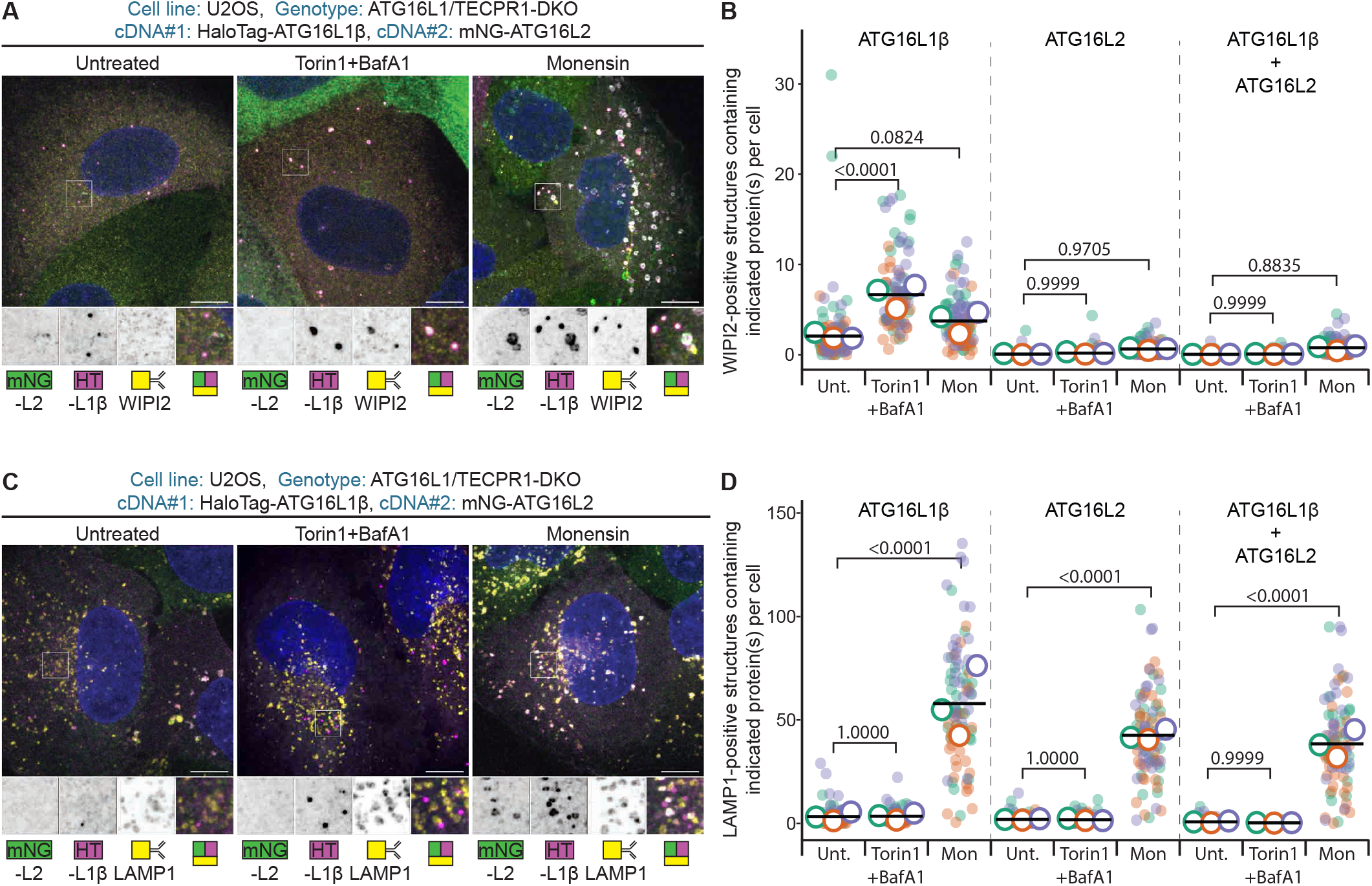
ATG16L2 is excluded from WIPI2 structures and recruited to compromised lysosomes. **A**. Confocal images of ATG16L1/TECPR1-DKO U2OS cells expressing HaloTag-ATG16L1β and mNeonGreen-ATG16L2, stained for WIPI2, left untreated or treated with 50 nM Torin1 plus 100 nM bafilomycin A1 for 30 min or 5 µM monensin for 15 min. Scale bars, 10 μm. **B**. Quantification of WIPI2-positive structures containing ATG16L1β, ATG16L2, or both proteins from images in (**A**). Total cells per condition/images per condition: 267/30 (Untreated), 289/30 (Torin1+BafA1), 285/30 (Monensin). **C**. Confocal images of ATG16L1/TECPR1-DKO U2OS cells expressing HaloTag-ATG16L1β and mNeonGreen-ATG16L2, stained for LAMP1, left untreated or treated with 50 nM Torin1 plus 100 nM bafilomycin A1 for 30 min or 5 µM monensin for 15 min. Scale bars, 10 μm. **D**. Quantification of LAMP1-positive structures containing ATG16L1β, ATG16L2, or both proteins from images in (**C**). Total cells per condition/images per condition: 335/30 (Untreated), 315/30 (Torin1+BafA1), 339/30 (monensin). **Data information:** Dot colors indicate independent experiments with each dot representing one image; large open symbols indicate the mean of each independent replicate; black horizontal lines indicate the overall mean. Statistical significance was determined by two-way ANOVA followed by Tukey’s multiple-comparisons test (**B,D**). Adjusted p-values are indicated in the graphs.

### The ATG16L2 CCD imposes a functional constraint on homodimer activity

To define the structural basis of ATG16L2 function, we used a previously validated chimera strategy in which the CCDs of ATG16L1β and ATG16L2 were exchanged (Ishibashi *et al*., 2011) (**Fig. 3A**). We generated untagged chimeras, in which the terminal domains of ATG16L1β or ATG16L2 were paired with the CCD of the other protein, termed 1-2-1 and 2-1-2 (where 1 = ATG16L1β and 2 = ATG16L2), respectively. Monensin-induced atg8ylation was assessed in the presence of VPS34 and ULK1 inhibitors to suppress canonical autophagy and thereby enrich for CASM-specific signal. This allowed us to test whether the otherwise inactive ATG16L2 terminal domains could support CASM when combined with the ATG16L1 CCD.

**Figure 3.**
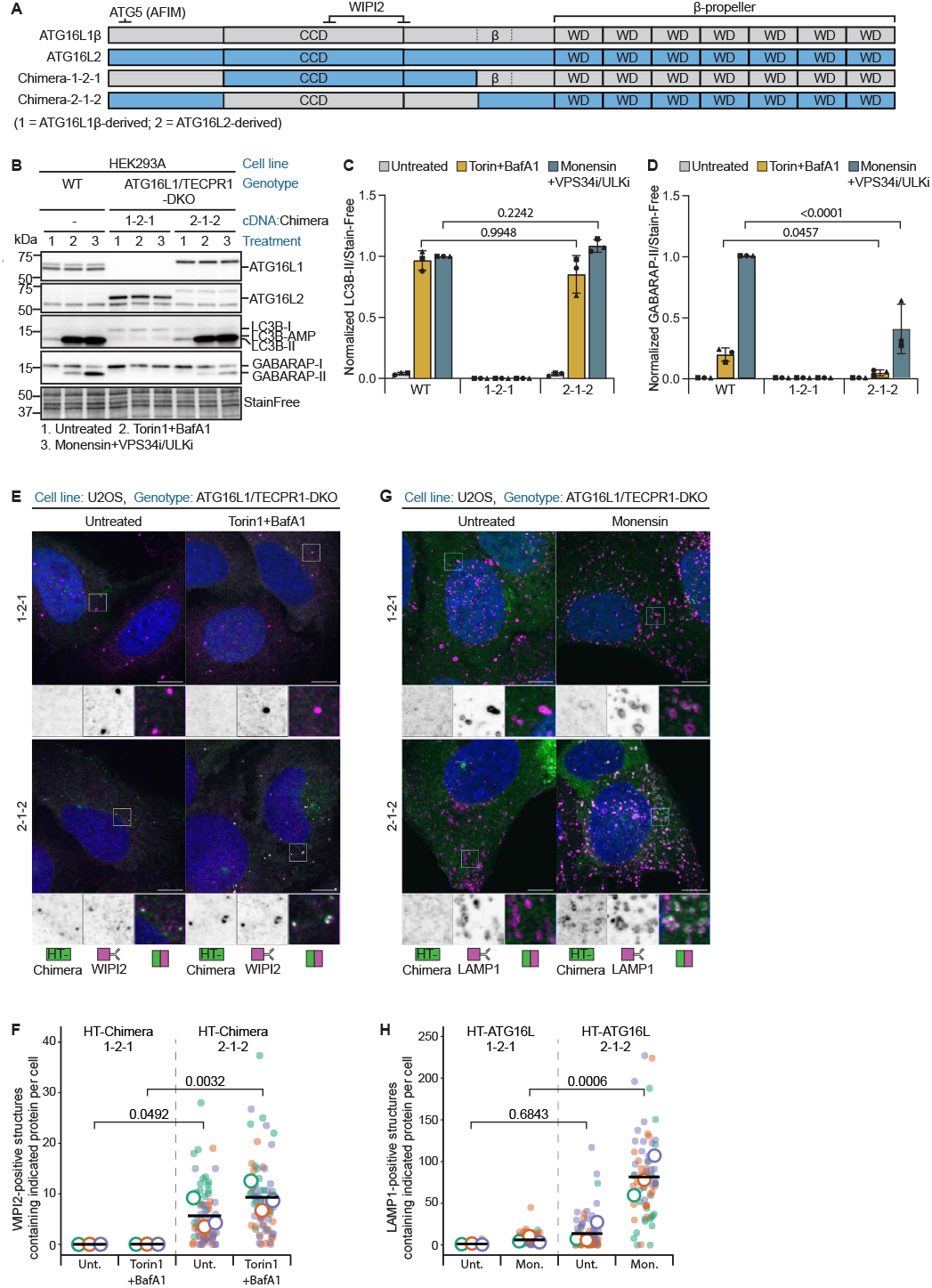
The ATG16L2 CCD imposes a functional constraint on homodimer activity. **A**. Schematic representation of ATG16L1β, ATG16L2 and the domain-swap chimeras 1-2-1 and 2-1-2. Numbers indicate the origin of the N-terminal/CCD/C-terminal regions, with 1 denoting ATG16L1β-derived sequence and 2 denoting ATG16L2-derived sequence. **B**. Immunoblot analysis of LC3B and GABARAP lipidation in WT and ATG16L1/TECPR1-DKO HEK293A cells expressing the indicated chimeras, left untreated (1), treated for 2 h with 50 nM Torin1 plus 100 nM bafilomycin A1 (2), or treated with 5 µM monensin together with 5 µM MRT68921 and 10 µM SAR405 (3). Stain-Free is a loading control. MRT68921 and SAR405 are ULK and VPS34 inhibitors, respectively. **C, D**. Quantification of LC3B-II (**C**) and GABARAP-II (**D**) from immunoblots in (**B**), normalized to Stain-Free. **E**. Confocal images of ATG16L1/TECPR1-DKO U2OS cells expressing HaloTag-1-2-1 or HaloTag-2-1-2, stained for WIPI2, left untreated or treated with 50 nM Torin1 plus 100 nM bafilomycin A1 for 30 min. Scale bars, 10 μm. **F**. Quantification of WIPI2-positive structures containing the indicated HaloTag chimera from images in (**E**). Total cells per condition/images per condition: 252/25 (1-2-1 untreated), 259/25 (1-2-1 Torin1+BafA1), 241/25 (2-1-2 untreated), 284/25 (2-1-2 Torin1+BafA1). **G**. Confocal images of ATG16L1/TECPR1-DKO U2OS cells expressing HaloTag-1-2-1 or HaloTag-2-1-2, stained for LAMP1, left untreated or treated with 5 µM monensin for 15 min. Scale bars, 10 μm. **H**. Quantification of LAMP1-positive structures containing the indicated HaloTag chimera from images in (**G**). Total cells per condition/images per condition: 244/25 (1-2-1 untreated), 248/25 (1-2-1 monensin), 236/25 (2-1-2 untreated), 222/25 (2-1-2 monensin). **Data information:** Quantifications are normalized as indicated in the graphs. Bars show mean ± SD; symbols denote independent experiments. Statistical significance was determined by two-way ANOVA followed by Tukey’s multiple-comparisons test (**C,D**). Adjusted p-values are indicated in the graphs. For image quantifications; Dot colors indicate independent experiments with each doth representing one image; large open symbols indicate the mean of each independent replicate; black horizontal lines indicate the overall mean. Statistical significance was determined by two-way ANOVA followed by Tukey’s multiple-comparisons test (**F,H**). Adjusted p-values are indicated in the graphs.

Consistent with previous work, the 2-1-2 chimera restored atg8ylation in DKO cells treated with Torin1 and bafilomycin A1, while the 1-2-1 chimera did not (**Fig. 3B–D**). Thus, ATG16L2 retains catalytic potential, but this is constrained by its native CCD, likely because this domain does not support productive homodimerization or WIPI2-dependent recruitment. Notably, 2-1-2 also supported GABARAP lipidation, indicating that the GABARAP lipidation defect seen in Fig. 1 is unlikely to reflect an intrinsic substrate preference of ATG16L2. Instead, it is more likely explained by the single-ATG5-binding-arm configuration of the functional ATG16L1-ATG16L2 heterodimer.

Under monensin stimulation in the presence of VPS34 and ULK1 inhibitors, 1-2-1 again remained inactive, whereas 2-1-2 was competent to drive atg8ylation (**Fig. 3B–D**). The inhibitory effect of the ATG16L2 CCD therefore extends to both canonical autophagy and CASM. Importantly, the ability of the ATG16L1 CCD to restore activity to the ATG16L2 terminal domains indicates that the CCD is a key determinant of whether the complex is functionally engaged.

In ATG16L1/TECPR1-DKO U2OS cells, HaloTag-2-1-2 was efficiently recruited to both WIPI2-positive autophagic structures and LAMP1-positive compartments under the respective inducing conditions. By contrast, HaloTag-1-2-1 failed to form Torin1-induced autophagic puncta, consistent with loss of productive WIPI2-dependent recruitment (**Fig. 3E–H**). HaloTag-1-2-1 also showed markedly reduced recruitment to LAMP1-positive compartments compared with HaloTag-2-1-2, indicating that the ATG16L2 CCD limits productive membrane engagement in both autophagy and CASM. Consistent with the untagged chimera, HaloTag-2-1-2 also restored atg8ylation in ATG16L1/TECPR1-DKO cells under both autophagy- and CASM-inducing conditions (**Figure EV3A–C**).

Together, these data show that the terminal domains of ATG16L2 are catalytically competent, but that this activity is constrained by its native CCD. Rather than supporting a functional complex on its own, ATG16L2 requires pairing with ATG16L1β, whose CCD is sufficient to confer catalytic competence.

### The β-propeller is a critical determinant of ATG16L2 function in VAIL

Because the ATG16L1β β-propeller is required for CASM, and an ATG16L2 chimera carrying the ATG16L1 CCD is CASM-competent, we asked whether the ATG16L2 β-propeller can functionally substitute for that of ATG16L1. To test this, we expressed mNeonGreen-tagged ATG16L2 either as the full-length protein or as a C-terminal truncation lacking the β-propeller (aa 1–321) in ATG16L1/TECPR1-DKO U2OS cells. Whereas full-length ATG16L2 was rapidly recruited to lysosomes after monensin treatment, the Δβ-propeller mutant remained cytosolic, showing that the β-propeller is required for recruitment to stressed lysosomes (**Fig. 4A,B**). These data further show that ATG16L2 can be recruited to lysosomes independently of ATG16L1β, even as a likely monomer without any atg8ylation activity.

**Figure 4.**
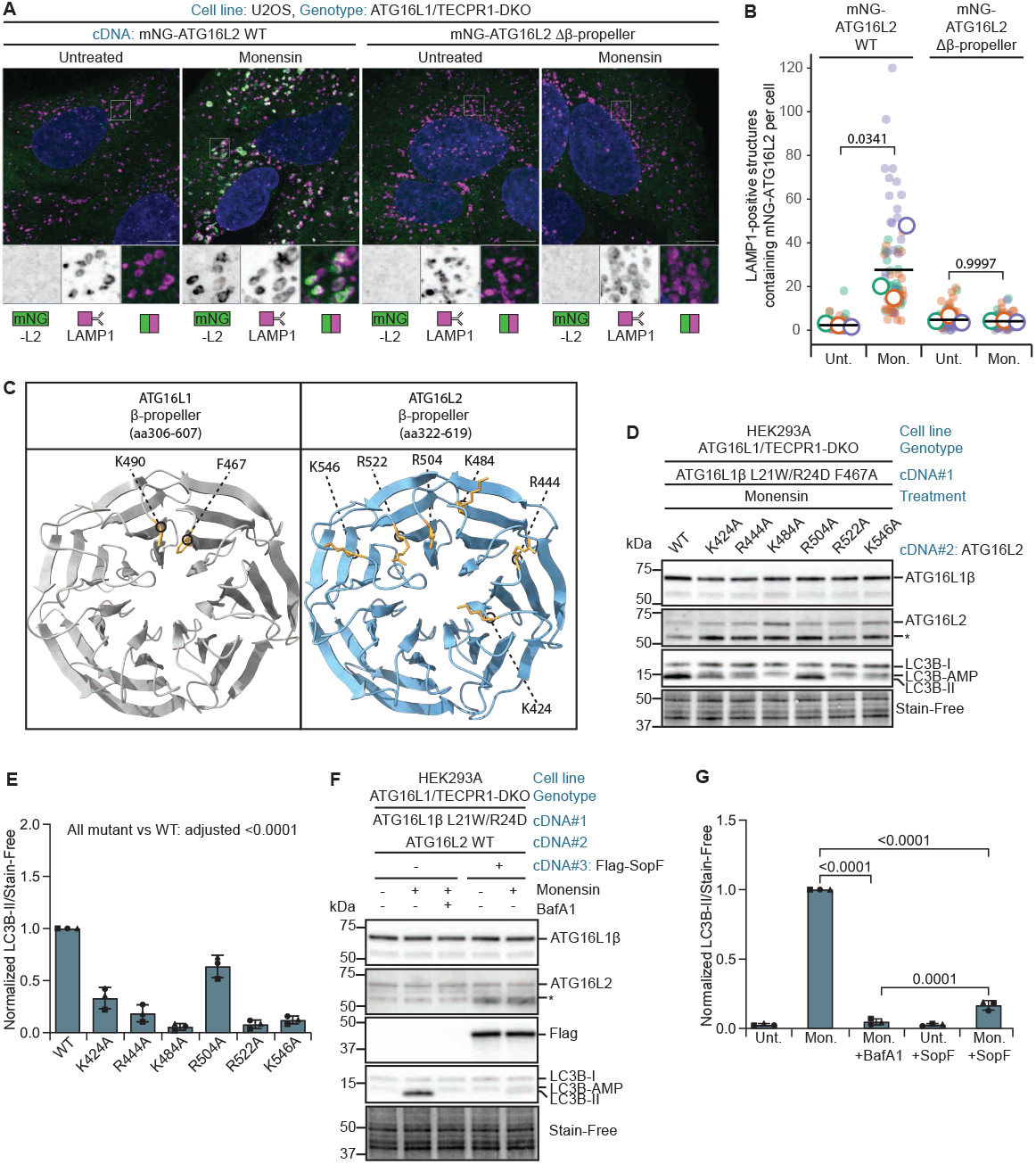
The β-propeller is a critical determinant of ATG16L2-dependent VAIL. **A**. Confocal images of ATG16L1/TECPR1-DKO U2OS cells expressing mNeonGreen-ATG16L2 WT or an ATG16L2 Δβ-propeller mutant, stained for LAMP1, left untreated or treated with 5 µM monensin for 15 min. Scale bars, 10 μm. **B**. Quantification of LAMP1-positive structures containing mNeonGreen-ATG16L2 from images in (**A**). Total cells per condition/images per condition: 284/30 (ATG16L2 untreated), 258/30 (ATG16L2 monensin), 245/30 (ATG16L2 Δβ-propeller untreated), 231/30 (ATG16L2 Δβ-propeller monensin). **C**. Structural comparison of the ATG16L1 and ATG16L2 β-propeller domains. The ATG16L1 β-propeller structure was obtained from the AlphaFold Protein Structure Database model AF-Q676U5-F1-v6 and visualized for residues aa306–607. The ATG16L2 β-propeller structure was obtained from the AlphaFold Protein Structure Database model AF-Q8NAA4-F1-v6 and visualized for residues aa322–619. Residues selected for alanine substitution in ATG16L2, and corresponding CASM-relevant residues in ATG16L1, are highlighted. **D**. Immunoblot analysis of LC3B lipidation in ATG16L1/TECPR1-DKO HEK293A cells expressing ATG16L1β L21W/R24D F467A together with WT or mutant ATG16L2 proteins, treated with 5 µM monensin for 2 h. **E**. Quantification of LC3B-II from immunoblots in (**D**), normalized to Stain-Free. All ATG16L2 mutants were compared with WT ATG16L2. **F**. Immunoblot analysis of LC3B lipidation in ATG16L1/TECPR1-DKO HEK293A cells expressing ATG16L1β L21W/R24D together with WT ATG16L2. Cells were left untreated, treated with 5 µM monensin, treated with 5 µM monensin plus 100 nM bafilomycin A1, or analyzed in the absence or presence of Flag-SopF as indicated. **G**. Quantification of LC3B-II from immunoblots in (**F**), normalized to Stain-Free. **Data information:** Quantifications are normalized as indicated in the graphs. Bars show mean ± SD; symbols denote independent experiments. Statistical significance was determined by one-way ANOVA followed by Tukey’s multiple-comparisons test (**E,G**). Adjusted p-values are indicated in the graphs. For image quantifications; Dot colors indicate independent experiments with each doth representing one image; large open symbols indicate the mean of each independent replicate; black horizontal lines indicate the overall mean. Statistical significance was determined by two-way ANOVA followed by Tukey’s multiple-comparisons test (**B**). Adjusted p-values are indicated in the graphs.

To identify functionally important residues within the ATG16L2 β-propeller, we structurally aligned AlphaFold-predicted ATG16L1β and ATG16L2 β-propeller models (Jumper *et al*, 2021; Varadi *et al*, 2022) using ChimeraX MatchMaker (Meng *et al*, 2006; Pettersen *et al*, 2021) and mutated positively charged ATG16L2 residues that mapped to the surface corresponding to ATG16L1β F467 and K490, two residues required for ATG16L1β-dependent VAIL (**Fig. 4C**) (Fletcher *et al*, 2018). We then tested these mutants in ATG16L1/TECPR1-DKO cells together with an ATG16L1β mutant defective in both CASM (F467A) and ATG5 binding (L21W, R24D) (**Fig. 4D,E**). Importantly, ATG16L1β-ATG16L2 heterodimers remained CASM-competent despite the VAIL-deficient F467A mutation in ATG16L1β, allowing us to isolate the contribution of the ATG16L2 β-propeller. Whereas R504A had only a minor effect, all other mutations impaired CASM, and K484A and R522A abolished activity completely (**Fig. 4D,E**). Interestingly, ATG16L2 K484 aligns with ATG16L1 K469 in the β-propeller, a residue recently implicated in VAIL regulation through deacetylation (Wang *et al*, 2025).

To determine whether the ATG16L1β-ATG16L2 heterodimer acts through the canonical VAIL pathway, we tested the effects of bafilomycin A1 and the *Salmonella* effector SopF. SopF inhibits CASM by ADP-ribosylating the V-ATPase subunit ATP6V0C, thereby preventing V-ATPase-dependent recruitment of ATG16L1 to stressed single membranes and blocking local ATG8 lipidation. Both bafilomycin A1 and SopF completely abolished atg8ylation (**Fig. 4F,G**), indicating that the heterodimer acts through the established V-ATPase-dependent pathway. Together, these findings identify ATG16L2 as a V-ATPase-responsive adaptor that mediates VAIL through a conserved β-propeller-dependent mechanism.

### ATG16L2 confers VAIL activity on the ATG16L1α isoform

Although both ATG16L1 splice variants support canonical macroautophagy, only the β-isoform is VAIL-competent, whereas the α-isoform is not (Lystad *et al*., 2019). This prompted us to ask whether ATG16L2 could act as a specialized binding partner that confers VAIL activity on ATG16L1α. To examine the localization of the ATG16L1 isoforms independently of endogenous ATG16L1, we expressed HaloTag-ATG16L1α or HaloTag-ATG16L1β in U2OS ATG16L1/TECPR1-DKO cells and monitored their recruitment during monensin-induced stress. ATG16L1α localized only to small WIPI2-positive autophagic structures (**Fig. 5A,B**) and not to LAMP1-positive compartments (**Fig. 5C,D**), consistent with an autophagy-restricted state. By contrast, ATG16L1β was recruited to both WIPI2- and LAMP1-positive structures, in line with its dual role in autophagy and CASM (**Fig. 5A–D**).

**Figure 5:**
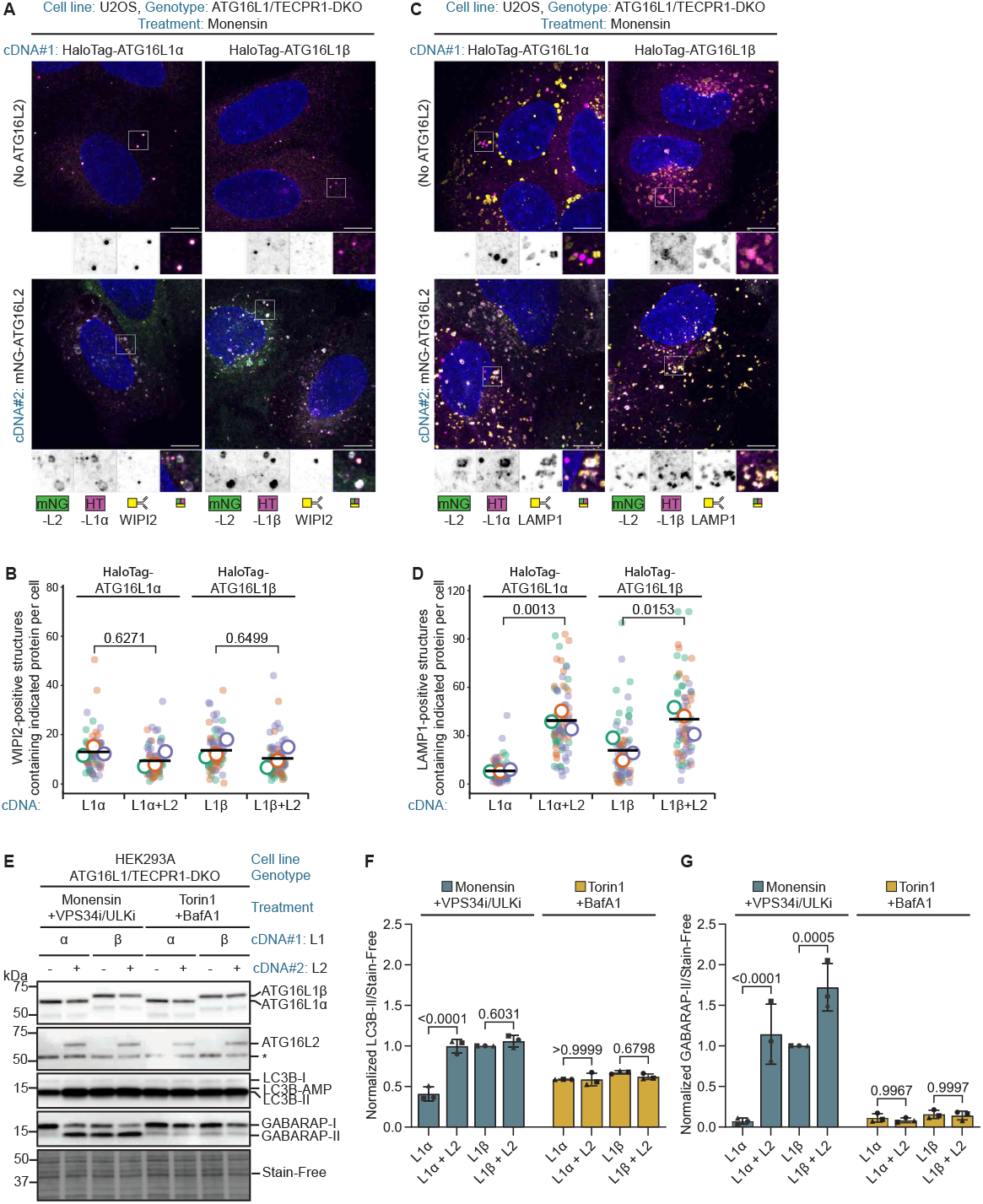
ATG16L2 enables ATG16L1α-dependent VAIL activity. **A**. Confocal images of ATG16L1/TECPR1-DKO U2OS cells expressing HaloTag-ATG16L1α or HaloTag-ATG16L1β in the absence or presence of mNeonGreen-ATG16L2, stained for WIPI2 after treatment with 5 µM monensin for 15 min. Scale bars, 10 μm. **B**. Quantification of WIPI2-positive structures containing the indicated HaloTag-ATG16L1 isoform from images in (**A**), in the absence or presence of mNeonGreen-ATG16L2. Total cells per condition/images per condition: 227/25 (ATG16L1α monensin), 261/25 (ATG16L1α + ATG16L2 monensin), 242/25 (ATG16L1β monensin), 232/25 (ATG16L1β + ATG16L2 monensin). **C**. Confocal images of ATG16L1/TECPR1-DKO U2OS cells expressing HaloTag-ATG16L1α or HaloTag-ATG16L1β in the absence or presence of mNeonGreen-ATG16L2, stained for LAMP1 after treatment with 5 µM monensin for 15 min. Scale bars, 10 μm. **D**. Quantification of LAMP1-positive structures containing the indicated HaloTag-ATG16L1 isoform from images in (**C**), in the absence or presence of mNeonGreen-ATG16L2. Total cells per condition/images per condition: 236/25 (ATG16L1α monensin), 241/25 (ATG16L1α + ATG16L2 monensin), 236/25 (ATG16L1β monensin), 263/25 (ATG16L1β + ATG16L2 monensin). **E**. Immunoblot analysis of LC3B and GABARAP lipidation in ATG16L1/TECPR1-DKO HEK293A cells expressing ATG16L1α or ATG16L1β in the absence or presence of ATG16L2, treated for 1 h with 5 µM monensin together with 5 µM MRT68921 and 10 µM SAR405 or with 50 nM Torin1 plus 100 nM bafilomycin A1. Stain-Free is a loading control. MRT68921 and SAR405 are ULK and VPS34 inhibitors, respectively. **F, G**. Quantification of LC3B-II (**F**) and GABARAP-II (**G**) from immunoblots in (**E**), normalized to Stain-Free. **Data information:** Quantifications are normalized as indicated in the graphs. Bars show mean ± SD; symbols denote independent experiments. Statistical significance was determined by two-way ANOVA followed by Tukey’s multiple-comparisons test (**F,G**). Adjusted p-values are indicated in the graphs. For image quantifications; Dot colors indicate independent experiments with each doth representing one image; large open symbols indicate the mean of each independent replicate; black horizontal lines indicate the overall mean. Statistical significance was determined by one-way ANOVA followed by Tukey’s multiple-comparisons test (**B,D**). Adjusted p-values are indicated in the graphs.

Strikingly, co-expression of mNeonGreen-ATG16L2 redistributed a substantial pool of ATG16L1α to LAMP1-positive lysosomal membranes, indicating that ATG16L2 converts ATG16L1α from an autophagy-restricted to a VAIL-competent state. Notably, ATG16L2 co-expression also enhanced recruitment of ATG16L1β to LAMP1-positive structures during CASM induction. In parallel, recruitment of both ATG16L1α and ATG16L1β to WIPI2-positive autophagic structures was modestly reduced, suggesting that ATG16L2 shifts the distribution of ATG16L1-containing complexes away from WIPI2-positive autophagic membranes and toward LAMP1-positive VAIL compartments.

We next asked whether this redistribution translates into increased atg8ylation. To suppress canonical autophagy-derived atg8ylation and thereby isolate the VAIL-specific response, cells were treated with monensin in the presence of VPS34 and ULK1/2 inhibitors. Under these conditions, ATG16L1/TECPR1-DKO HEK293A cells reconstituted with ATG16L1α showed minimal atg8ylation, whereas ATG16L1β supported robust LC3B and GABARAP lipidation (**Fig. 5E–G**). Notably, co-expression of ATG16L2 with ATG16L1α restored VAIL activity to levels comparable to ATG16L1β. In contrast to the engineered single-arm ATG16L1β mutant–ATG16L2 complex described above (**Fig. 1H, Figure EV1D,E**), this native ATG16L1α–ATG16L2 heterodimer contained two functional ATG5-binding arms and supported efficient lipidation of both LC3B and GABARAP.

Together with the AFIM-mutant rescue experiments, these findings suggest that one functional ATG5-binding arm is sufficient to support partial LC3B lipidation during VAIL, whereas robust GABARAP lipidation requires a bivalent ATG16L1α/β–ATG16L2 dimer in which both arms can bind ATG5. Consistent with this model, ATG16L2 also enhanced GABARAP lipidation in ATG16L1β-expressing cells, indicating that ATG16L2 can expand the pool of VAIL-dedicated E3-like heterodimers even when the VAIL-competent β-isoform is present. In contrast, LC3B lipidation was not further increased by ATG16L2 co-expression, possibly because ATG16L1β already supports near-maximum levels of LC3B lipidation under these conditions.

Together, these data identify the ATG16L1α/β-ATG16L2 heterodimer as a specialized V-ATPase-responsive complex that enables robust VAIL by overcoming the recruitment constraints of ATG16L1α and improving the recruitment response of both ATG16L1 isoforms.

## Discussion

Our data identify ATG16L2 as a dedicated VAIL factor whose activity is unlocked by heterodimerization with ATG16L1. This provides a mechanistic explanation for an unresolved paradox in ATG16L2 biology: despite being a close ATG16L1 paralogue that can assemble with ATG12–ATG5, ATG16L2 does not support canonical autophagy (Ishibashi *et al*., 2011). Earlier chimera studies suggested that this functional divergence is largely determined by the CCD (Ishibashi *et al*., 2011). Our results extend this model by showing that the ATG16L2 CCD does not merely explain a lack of canonical autophagy activity, but instead constrains ATG16L2 in a latent state that becomes functional only upon heterodimerization with ATG16L1α/β.

These findings further distinguish canonical autophagy and CASM at the level of ATG16 complex composition and membrane targeting. In canonical autophagy, ATG16L1 is recruited to PtdIns(3)P-positive phagophores through WIPI2 (Dooley *et al*., 2014). By contrast, CASM depends on the C-terminal β-propeller of ATG16L1 and proceeds through a V-ATPase–ATG16L1 axis that is sensitive to bafilomycin A1 and inhibited by SopF (Fischer *et al*, 2020; Fletcher *et al*., 2018; Xu *et al*, 2019).

Our data place ATG16L2 in this V-ATPase-dependent branch. ATG16L2 is recruited to stressed lysosomes, requires its own β-propeller for VAIL activity, and shifts its associated ATG16L1 molecule away from WIPI2-positive autophagic structures. Thus, ATG16L1–ATG16L2 heterodimers define a VAIL-selective form of the ATG16 machinery. In this configuration, ATG16 complex composition helps determine whether ATG16L1 engages WIPI2-dependent canonical autophagy or is redirected toward V-ATPase-responsive atg8ylation.

A central conclusion from our study is that this specialization is imposed by the ATG16L2 CCD. Rather than serving only as a dimerization interface, this region restricts productive ATG16L2 complex formation. Replacing it with the ATG16L1 CCD was sufficient to restore both membrane recruitment and atg8ylation, showing that the terminal domains of ATG16L2 are functionally competent once this constraint is relieved. We therefore propose that ATG16L2 acts as a molecular gatekeeper that channels ATG16L1 into a VAIL-dedicated heterodimeric complex. This model is most consistent with ATG16L2 competing with ATG16L1 during de novo complex assembly, rather than replacing ATG16L1 within pre-existing homodimers. Although our data does not directly resolve the timing of heterodimer formation, they show that ATG16L2 shifts the steady-state pool of ATG16L1-containing complexes toward VAIL competence. Such specialization could allow cells to preserve rapid VAIL-type single-membrane atg8ylation even when the canonical autophagy machinery is engaged elsewhere.

This model is supported by the ability of ATG16L2 to confer VAIL activity on ATG16L1α. We have previously shown that both ATG16L1 isoforms support canonical autophagy, whereas only ATG16L1β efficiently supports VAIL. Here, we show that ATG16L2 overcomes this limitation by redirecting ATG16L1α from an autophagy-restricted state into a lysosome-associated, VAIL-competent heterodimer. Notably, this native heterodimer restored lipidation across ATG8 subfamilies, unlike an engineered complex with only one functional ATG5-binding arm, which supported LC3B but not GABARAP conjugation. Thus, ATG16L2 expands ATG16L1α function by generating a distinct, fully competent VAIL E3-like complex.

Transcript diversity should also be considered when interpreting ATG16L2 function. As noted above, we focused on full-length human ATG16L2, which contains the N-terminal ATG5-interaction motif, central CCD and C-terminal WD40 β-propeller required for ATG5 engagement, ATG16L1 heterodimerization and lysosomal recruitment. Thus, our data define the activity of full-length ATG16L2-containing heterodimers, while leaving possible roles for shorter or tissue-restricted ATG16L2 transcripts unresolved.

The low abundance of ATG16L2 in standard laboratory cell lines may have obscured elucidation of its physiological role. Our data raise the possibility that ATG16L2 is deployed in selected cell types or activation states where single-membrane stress is particularly prominent. Myeloid cells are an attractive context in which to consider this possibility, as high phagocytic load, membrane damage and lysosomal stress may create a need for a dedicated VAIL-responsive ATG16 complex.

Together, our findings identify ATG16L2 as a selective organizer of VAIL-competent ATG16 complexes. By heterodimerizing with ATG16L1α and β, ATG16L2 redirects ATG16L1 away from WIPI2-positive autophagic membranes and toward V-ATPase-responsive single-membrane compartments. A structural explanation for this shift may come from the dual WIPI2-binding mode of ATG16L1. Gong et al. showed that ATG16L1 engages WIPI2 through two sites, WBS1 and WBS2 (**Fig. 1A**), both of which are required for efficient autophagic flux (Gong *et al*., 2023). Whereas WBS1 forms a discrete binding element, WBS2 is embedded in the dimeric ATG16L1 coiled-coil and includes contributions from both ATG16L1 chains. In an ATG16L1α/β–ATG16L2 heterodimer, the non-conserved ATG16L2 coiled-coil surface would therefore be expected to disrupt the composite WBS2 interface.

This provides a mechanism by which cells can generate distinct ATG16 assemblies with different membrane specificities: ATG16L1α homodimers selectively support WIPI2-dependent canonical autophagy, whereas ATG16L1α/β–ATG16L2 heterodimers form a VAIL-dedicated branch of the ATG8 conjugation machinery. More broadly, these data show that diversification of ATG16 complex composition can direct the shared ATG8 conjugation machinery toward distinct membrane-remodeling pathways.

## Materials and Methods

### Plasmid constructs

Plasmid constructs are listed in Table EV1 and construction details are available upon request. All plasmids were verified by whole plasmid sequencing (Eurofins genomics).

### Cell culture

U2OS cells (ATCC, HTB-96) and HEK293A cells (a gift from Tamatsu Yoshimori, Osaka University) were maintained in Dulbecco’s Modified Eagle Medium (DMEM; Sigma-Aldrich, D0819) supplemented with 10% fetal cell serum (FCS; Sigma-Aldrich, F7524), 5 U/ml penicillin, and 50 μg/ml streptomycin. Cells were maintained at 37 °C in a humidified atmosphere with 5% CO_2_.

### Immunoblotting antibodies

ATG16L1 mouse mAb (MBL, M150-3, 1:1,000), ATG16L2 rabbit pAb (Proteintech, 24322-1-AP, 1:1000), ATG5 rabbit pAb (CST, 2630, 1:1000), DYKDDDDK (FLAG) rabbit mAb (CST, 14793, 1:1000), GABARAP (E1J4E) rabbit mAb (CST, 13733, 1:1,000), LC3B (D11) XP rabbit mAb (CST, 3868, 1:1,000), Halo Tag®mouse mAb (Promega, G9211, 1:1000), V5 mouse mAb (Abcam, ab27671, 1:1000), goat anti-mouse (Jackson ImmunoResearch, 115-035-003, 1:5,000) and HRP goat anti-rabbit (Jackson ImmunoResearch, 111-035-144, 1:5,000).

### Immunofluorescence and antibodies

Cells were seeded in Ibidi µ-Slide 18 Well dishes (Ibidi, 81816), stained with HaloTag ligand where indicated, treated as described, and fixed in 4% EM-grade formaldehyde in PBS (Cytiva Life Sciences, SH30256.01) for 12 min at 37°C. Cells were washed three times in PBS and permeabilized/blocked in PBS containing 0.05% saponin and 3% BSA for 20 min at room temperature. Cells were then incubated with the indicated primary antibodies diluted in PBS containing 3% BSA for 1 h at room temperature. After three washes in PBS containing 0.05% saponin, cells were incubated with secondary antibodies diluted in PBS containing 3% BSA and 10 µg/ml Hoechst 33342 (Thermo Scientific, H3570) for 45 minutes at RT. Cells were washed three times with PBS containing 0.05% saponin and once with PBS before imaging.

LAMP1 (clone H4A3) mouse mAb (H4A3 was deposited to the DSHB by August, J.T./Hildreth, J.E.K. (DSHB Hybridoma Product H4A3), 1:200, WIPI2 [2A2], Mouse, mAb (Abcam, ab105459) 1:200, Alexa488 donkey anti-mouse (Jackson ImmunoResearch, 715-545-150, 1:500), Alexa568 donkey anti-rabbit (Jackson ImmunoResearch, 711-575-152, 1:500), Alexa647 donkey anti-mouse (Jackson ImmunoResearch, 715-605-150, 1:500), Janelia Fluor® 549 HaloTag® Ligand (Promega, GA1110), Janelia Fluor® 646 HaloTag® Ligand (Promega, GA1120)

### Reagents

Reagents used in this study were: astemizole (Sigma-Aldrich, A2861), bafilomycin A1 (Enzo Life Sciences, BML-CM110), Buffer BXT 10× Strep-Tactin XT elution buffer (IBA Lifesciences, 2-1042-025), cOmplete Mini EDTA-free Protease Inhibitor Cocktail (Roche, 11836170001), Criterion 4–20% gradient 18-well Tris–HCl gels (Bio-Rad, 345-0034), DMEM with GlutaMAX (Gibco, 61965-026), doxycycline hyclate (Sigma-Aldrich, D9891), DTT (Sigma-Aldrich, D0632), fetal bovine serum (FBS; Sigma-Aldrich, F7524), Gly-Phe β-naphthylamide (GPN; Abcam, ab145914), H-Leu-Leu-OMe hydrochloride (LLOMe; Santa Cruz Biotechnology, SC-285992), monensin sodium salt (Sigma-Aldrich, M5273), NH_4_Cl (Sigma-Aldrich, A9434), nigericin sodium salt (Sigma-Aldrich, N7143), Hyclone Phosphate Buffered Saline (Cytiva Life Sciences, SH30256.01), Strep-Tactin XT magnetic beads (IBA Lifesciences, 2-4090-010), and Torin 1 (Bio-Techne R&D Systems, 4247).

### Production of stable lentiviral cell lines

Third-generation lentivirus were generated using procedures and plasmids as previously described (Campeau *et al*, 2009). Briefly, tagged fusions of transgenes were generated as Gateway ENTRY constructs using standard molecular biology techniques. From these vectors, lentiviral transfer vectors were generated by Gateway LR recombination into lentiviral destination vectors (Gateway-enabled vectors); pCDH-PGK-IRES-BLAST-DEST (System Biosciences) in addition to pLenti-PGK-Puro-DEST (Addgene plasmid 19068, a gift from Eric Campeau & Paul Kaufman). VSV-G pseudotyped lentiviral particles were packaged using a third-generation packaging system (Addgene plasmids 12251, 12253, and 12259, a gift from Didier Trono). Cells were then transduced with low virus titers and stable expressing populations were generated by antibiotic selection.

### Fluorescence-Activated Cell Sorting (FACS) of cell lines with fluorescently labeled proteins

To ensure comparable expression levels of fluorescently tagged proteins across cell lines, transduced populations were sorted by fluorescence-activated cell sorting (FACS). Cells were resuspended in complete medium to a concentration of 5-10 million cells/ml and passed through a 40 μm cell strainer prior to sorting. Fluorescent signals were detected using appropriate laser and filter settings, and gating was performed to exclude debris and doublets based on forward and side scatter parameters. Sorted cells were collected into complete growth medium and expanded prior to downstream assays.

### Stable transgenesis of HEK293A cells by piggyBac (PB) transposons

To generate cell lines stably expressing 3XFLAG-SopF, the cDNA encoding 3XFLAG-SopF was inserted into the PB-TA-ERN destination vector (Addgene plasmid #80474; a gift from Knut Woltjen) via Gateway LR recombination. The resulting PB-TA-ERN-3XFLAG-SopF expression vector was co-transfected with the Super PiggyBac Transposase expression vector into the cell lines indicated in figure 4F using FuGENE HD (Promega). Following transfection, PB-transgenic cells were successfully isolated through selection with 2 μg/ml puromycin.

### Generation of knockout cells using CRISPR/Cas9

Knockout cells were generated using the pX459 system (Ran *et al*, 2013). In short, cells were transfected using FuGENE HD (Promega) with pX459 vector (Addgene plasmid 62988, a gift from Feng Zhang) containing guides against ATG16L1 or TECPR1. Twenty-four hours post-transfection, transfected cells were selected with 2 μg/ml puromycin for 72 h. The population of cells that survived selection was then diluted and seeded to obtain clones originating from single cells. The individual clones were expanded before knockouts were confirmed by western blotting. Guide sequence for TECPR1 were from (Wible *et al*, 2019), while the guides used for ATG16L1 was adapted from our previously published work (Lystad *et al*., 2019), guide sequences are included in Table EV1.

### Western blotting

Cells were washed with ice-cold PBS and harvested in Triton X-100 lysis buffer (50 mM Tris–HCl pH 7.4, 150 mM NaCl, 1 mM EDTA, and 1% Triton X-100) supplemented with protease and phosphatase inhibitor cocktails and 40 mM N-ethylmaleimide (NEM). Lysates were incubated at 4°C for 30 min with end-over-end rotation, followed by centrifugation at 21,000 *g* for 10 min at 4°C to remove cellular debris. Protein concentrations of the cleared extracts were determined using the Pierce BCA protein assay (Thermo Fisher Scientific) and normalized with lysis buffer.

Samples were prepared by mixing extracts with Laemmli Sample Buffer containing dithiothreitol (DTT) and denatured at 95°C for 5 min. Equal amounts of protein (10 μg per lane) were resolved on Criterion 4–20% gradient TGX Stain-Free precast gels (Bio-Rad), activated for 30 seconds using the ChemiDoc™ MP Imaging System (Bio-Rad) and then transferred to low-fluorescence (LF) PVDF membranes using the Trans-Blot Turbo Rapid Transfer system (Bio-Rad) before being imaged for Stain-Free signals using the ChemiDoc™ MP Imaging System.

Membranes were blocked with EveryBlot Blocking Buffer (Bio-Rad) for 5 min at room temperature (RT), before overnight incubation with primary antibodies (diluted in 5% BSA in PBS-T [0.05% Tween-20]) at 4°C. Following primary incubation, membranes were washed and incubated with HRP-conjugated anti-rabbit or anti-mouse secondary antibodies in EveryBlot Blocking Buffer for 45 minutes at RT.

Protein bands were visualized using SuperSignal West Dura Extended Duration Substrate (Thermo Fisher Scientific) and imaged on an Azure Sapphire Biomolecular Imager. Densitometric analysis of protein bands was performed using AzureSpot (v2.1.097), while Stain-Free signals were quantified using Image Lab software (v6.1, Bio-Rad).

### Strep-TactinXT pull-down

Cells were trypsinized, resuspended in complete medium and counted. For each condition, 3 × 10^7^ cells were collected by centrifugation at 350 × g for 5 min, washed once in PBS, and the PBS was aspirated completely. Cell pellets were lysed in 600 µl Triton X-100 lysis buffer [50 mM Tris–HCl pH 7.4, 150 mM NaCl, 1 mM EDTA and 1% Triton X-100, supplemented with protease inhibitors] and incubated end-over-end for 30 min at 4°C. Lysates were cleared by centrifugation at 21,000 × g for 10 min at 4°C, and protein concentrations were determined by BCA assay. Cleared lysates were adjusted to equal protein concentrations before pull-down.

For each sample, 30 µl lysate was retained as input, and the remaining lysate was incubated with 100 µl pre-washed MagStrep Strep-TactinXT beads for 15 min at 4°C with shaking at 1,400 rpm. Beads were washed four times with 500 µl lysis buffer, with 1 min incubation at 4°C and 1,400 rpm for each wash. After the final wash, residual buffer was removed completely.

Bound proteins were eluted directly from the beads by addition of 60 µl sample buffer composed of 6 parts 2X Bio-Rad sample buffer, 1 part Strep-TactinXT elution buffer, 1 part 1 M DTT and 2 parts water. Input samples were prepared by mixing 10 µl input lysate with 20 µl sample buffer. For immunoblot analysis, 30 µl input sample and 20 µl pull-down sample were separated by SDS–PAGE and transferred to individual membranes for probing with the indicated antibodies.

### Microscopy

The fixed samples were imaged on a Nikon ECLIPSE Ti2-E inverted microscope (Nikon Corp, Tokyo, Japan) equipped with a Plan Apo λ 100× (NA 1.45, Oil) objective, a CSU-W1 SoRa spinning disk confocal unit (Yokogawa Electric Corp., Tokyo, Japan) equipped with a 50 μm pinhole disk and operated in SoRa mode using 1× magnification, two Prime BSI sCMOS cameras (Teledyne Photometrics, Tucson, AZ, US), a laser unit with 405/488/561/638 nm lasers (120/100/100/100 mW), a multichannel LED light source (Lumencor SprectraX Chroma, Lumencor, OR, US) and with QUAD (DAPI/GFP/RFP/Af647) filter cubes for epifluorescence. Images were processed into Maximum Projections and exported to TIF in NIS-Elements AR Analysis software, and processed into figure panels using the imageJ plugin FigureJ (Zinck & Mutterer, 2013). Image analysis was performed in CellProfiler (see below).

Multichannel images (DAPI/GFP/RFP/AF647 or DAPI/GFP/AF647) of 25-30 random fields of view (FOV) from each coverslip were captured. For each FOV, 21 sections were acquired (Z section spacing 0.4 μm). Images shown are maximum intensity projections of Z-sections and were adjusted by linear brightness-contrast adjustments.

### Image analysis

CellProfiler 4.2.6 software (Stirling *et al*, 2021) was used to identify and segment nuclei, cells and puncta to determine the per-cell number of puncta of different structures (ATG16L1α, ATG16L1β,ATG16L2, ATG16L2 β-propeller, ATG16L 1-2-1 and 2-1-2 chimeras, LAMP1, WIPI2 or LC3B) and their co-occurrence. The representative panel images were processed and generated using ImageJ (Fiji) (Schindelin *et al*, 2012).

### Statistics and reproducibility

Western blot data were analyzed and plotted using GraphPad Prism version 10.4.1. Microscopy data were analyzed and plotted in R (version 4.6.0) and RStudio (version 2025.09.2+418). The number of independent experiments, as well as the number of cells analyzed for microscopy, are indicated in the corresponding figure legends. Statistical tests used for each experiment are specified in the figure legends, and exact *P* values are reported. In instances of multiple comparisons of means, we used one-way or two-way analysis of variance (ANOVA) followed by Tukey’s to determine statistical significance, unless specified otherwise.

## Acknowledgements

The Core Facilities for Advanced Light Microscopy and Advanced Electron Microscopy at Oslo University Hospital and the MolMed Imaging Platform (MIP) at the Institute of Basic Medical Sciences, University of Oslo, are acknowledged for providing access to and training on relevant microscopes. NK and AHL were supported by a Young Research Talents Grant from the Research Council of Norway (Project number 325305). HS was supported by grants from the Norwegian Cancer Society (project number 182698), the South-Eastern Norway Regional Health Authority (project number 2016087), and the Research Council of Norway (project number 302994). This work was partly supported by the Research Council of Norway through its Centres of Excellence funding scheme (project number 262652). Figures were created using Adobe Illustrator CS6.

## Expanded View Figure Legends

**Figure EV1.**
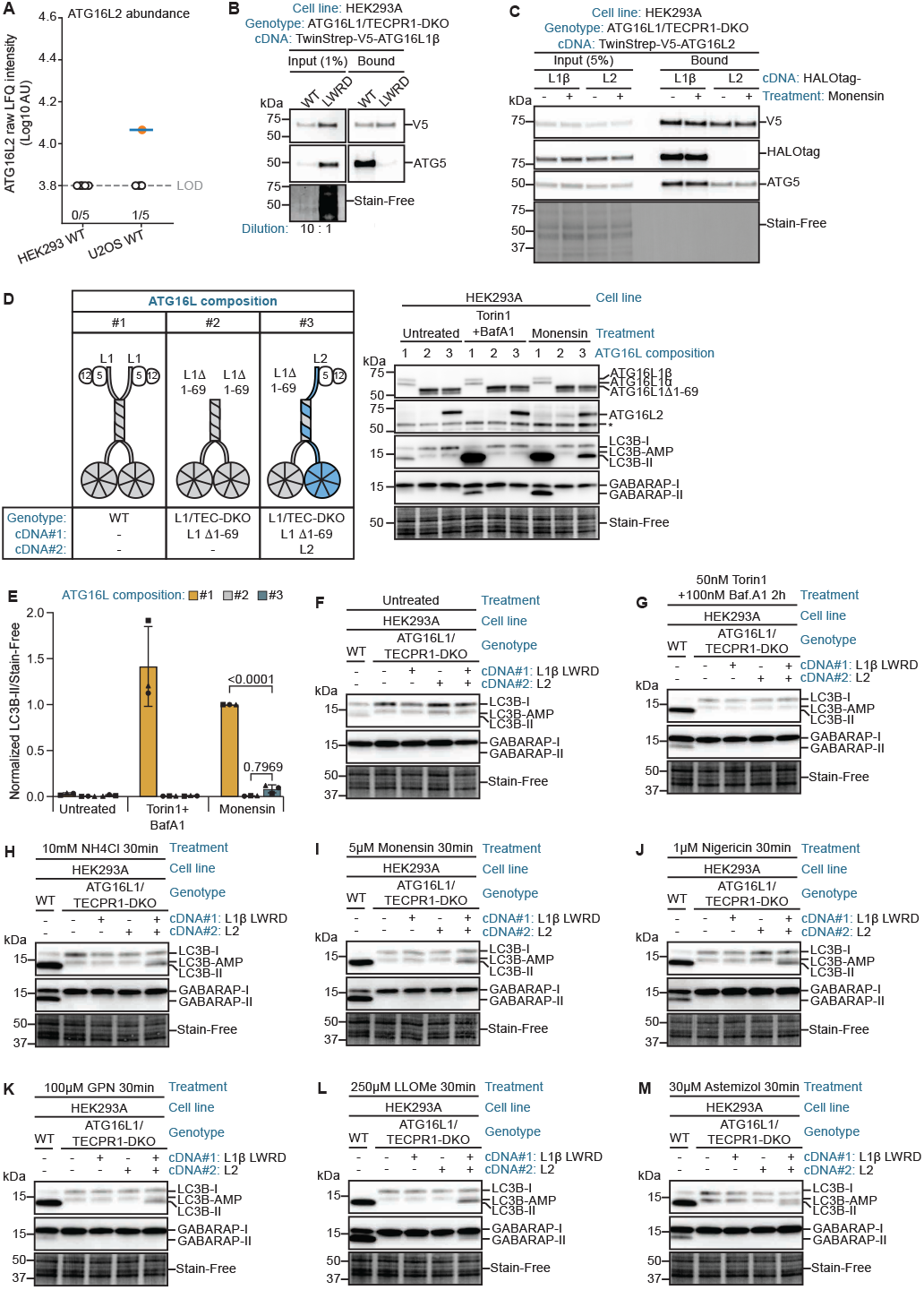
ATG16L2 expression and heterodimer activity across VAIL-inducing conditions. **A**. MS-based abundance of ATG16L2 in WT HEK293A and U2OS cells. Raw LFQ intensities are shown relative to the limit of detection (LOD). **B**. Strep-TactinXT pull-down of TwinStrep-V5-ATG16L1β WT or LWRD mutant from ATG16L1/TECPR1-DKO HEK293A cells, showing ATG5 binding in the input and pull-down fractions. For the WT sample, a 10-fold diluted lysate was used, resulting in 10-fold less protein loaded in the input and 10-fold less material incubated with the Strep-TactinXT beads, as indicated. **C**. Strep-TactinXT pull-down of TwinStrep-V5-ATG16L2 from ATG16L1/TECPR1-DKO HEK293A cells co-expressing HaloTag-ATG16L1β or HaloTag-ATG16L2, left untreated or treated with 5 µM monensin for 30 min. **D**. Schematic of the ATG16L compositions analyzed and corresponding immunoblot of LC3B and GABARAP lipidation in HEK293A cells expressing the indicated ATG16L combinations, left untreated or treated with 50 nM Torin1 plus 100 nM bafilomycin A1 or 5 µM monensin for 2 h. ATG16L composition #1, WT cells expressing endogenous ATG16L1; ATG16L composition #2, ATG16L1/TECPR1-DKO cells expressing ATG16L1β aa70-607; ATG16L composition #3, ATG16L1/TECPR1-DKO cells co-expressing ATG16L1β aa70-607 and ATG16L2. Stain-Free is a loading control. **E**. Quantification of LC3B-II from immunoblots in (**D**), normalized to Stain-Free. **F–M**. Immunoblot analysis of LC3B and GABARAP lipidation in WT and ATG16L1/TECPR1-DKO HEK293A cells xpressing ATG16L1β L21W/R24D with or without ATG16L2, treated as indicated. Stain-Free is a loading control.

**Figure EV2.**
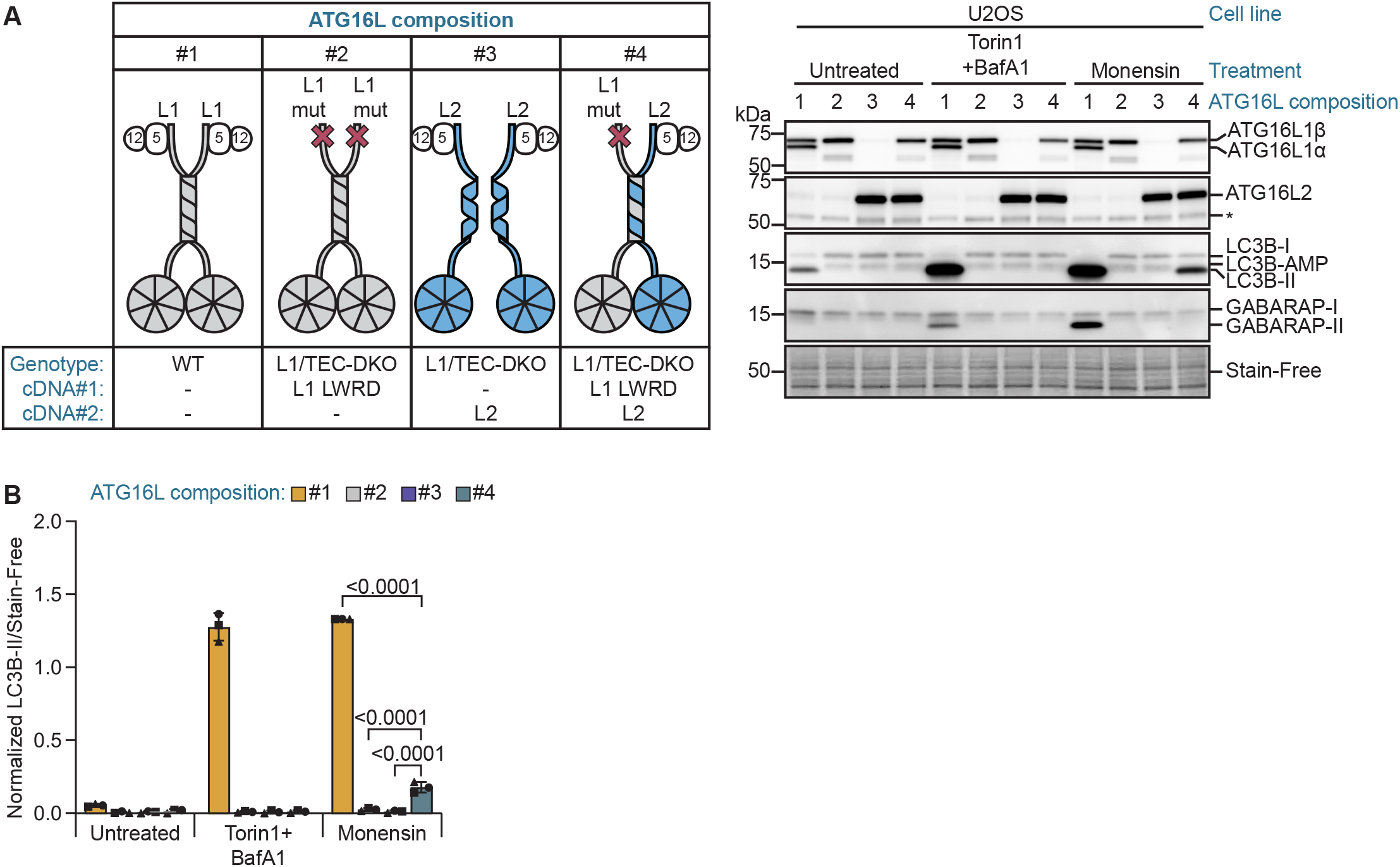
ATG16L2 requires heterodimerization with ATG16L1 for atg8ylation in U2OS cells. **A**. Schematic of the ATG16L compositions analyzed and corresponding immunoblot of LC3B and GABARAP lipidation in U2OS cells expressing the indicated ATG16L combinations, left untreated or treated with 50 nM Torin1 plus 100 nM bafilomycin A1 or 5 µM monensin for 2 h. ATG16L composition #1, WT cells expressing endogenous ATG16L1; ATG16L composition #2, ATG16L1/TECPR1-DKO cells expressing ATG16L1β L21W/R24D; ATG16L composition #3, ATG16L1/TECPR1-DKO cells expressing ATG16L2; composition #4, ATG16L1/TECPR1-DKO cells co-expressing ATG16L1β L21W/R24D and ATG16L2. Stain-Free is a loading control. **B** Quantification of LC3B-II from immunoblots in (**A**), normalized to Stain-Free. **Data information:** Quantifications are normalized as indicated in the graphs. Bars show mean ± SD; symbols denote independent experiments. Statistical significance was determined by two-way ANOVA followed by Tukey’s multiple-comparisons test (**B**). Adjusted p-values are indicated in the graphs.

**Figure EV3.**
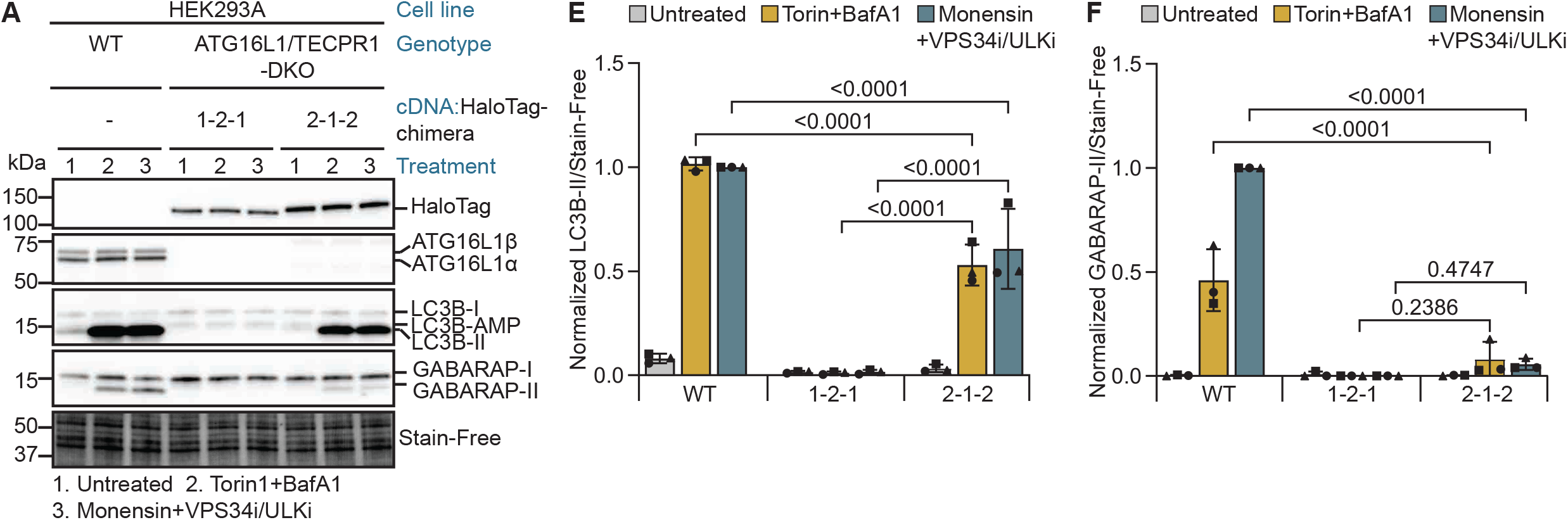
The HaloTag-2-1-2 chimera restores atg8ylation in HEK293A cells during autophagy and VAIL. **A**. Immunoblot analysis of LC3B and GABARAP lipidation in WT and ATG16L1/TECPR1-DKO HEK293A cells expressing the indicated chimeras, left untreated (1), treated for 2 h with 50 nM Torin1 plus 100 nM bafilomycin A1 (2), or treated with 5 µM monensin together with 5 µM MRT68921 and 10 µM SAR405 (3). Stain-Free is a loading control. MRT68921 and SAR405 are ULK and VPS34 inhibitors, respectively. **B, C**. Quantification of LC3B-II (**B**) and GABARAP-II (**C**) from immunoblots in (**A**), normalized to Stain-Free. **Data information:** Quantifications are normalized as indicated in the graphs. Bars show mean ± SD; symbols denote independent experiments. Statistical significance was determined by two-way ANOVA followed by Tukey’s multiple-comparisons test (**B,C**). Adjusted p-values are indicated in the graphs.

**EV_Table_1:**
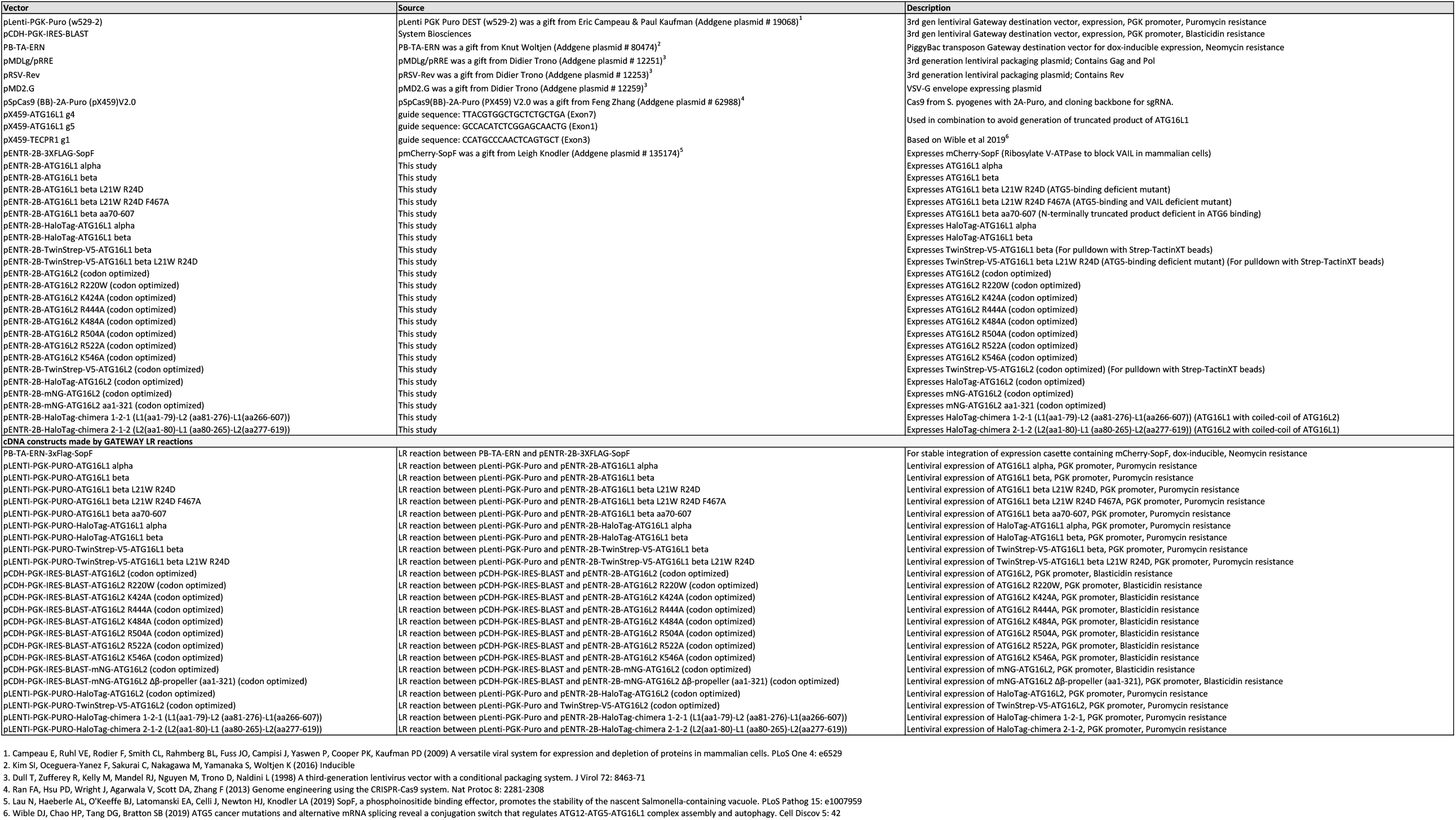
Constructs used in this study.

